# MoveR: an R package for easy processing and analysis of animal video-tracking data

**DOI:** 10.1101/2023.11.08.566216

**Authors:** Quentin Petitjean, Silène Lartigue, Mélina Cointe, Nicolas Ris, Vincent Calcagno

## Abstract

Animal movement and behavior are critical to understanding ecological and evolutionary processes. Recent years have witnessed an increase in methodological and technological innovations in video-tracking solutions for phenotyping animal behavior. Although these advances enable the collection of high-resolution data describing the movement of multiple individuals, analyzing and interpreting them remains challenging due to their complexity, heterogeneity, and noisiness. Here, we introduce MoveR, an R package for importing, filtering, visualizing, and analyzing data from common video-tracking solutions. MoveR includes flexible tools for polishing data, removing tracking artifacts, subsetting and plotting individual paths, and computing different movement and behavior metrics.

**Metadata:** 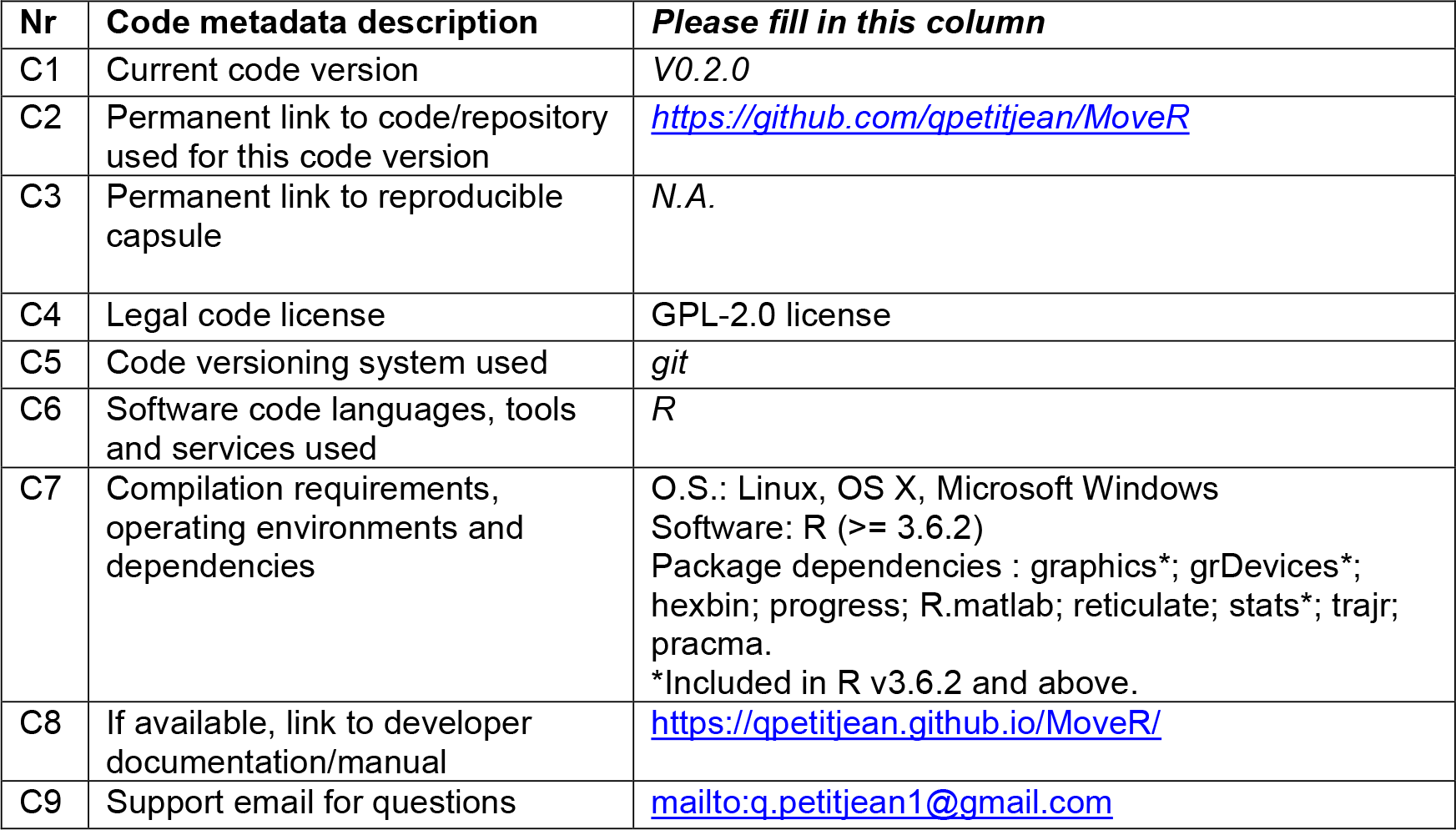

## 1. Motivation and significance

### 1.1. Scientific motivation

Studying animal movement patterns and behaviors is paramount to understanding ecological and evolutionary processes and developing effective species management strategies. Animal movements and behavior are often considered the first response to environmental changes or stressful conditions (e.g., pollution, climate change, introduction of non-native species) [1,2], making them valuable tools to infer underlying animal’s internal states, sensory biases, and behavioral types. Therefore, changes in movement patterns and behavioral traits are increasingly considered a highly integrative and relevant endpoint to assess the effect of global changes and anthropogenic stress on organisms (e.g., ecotoxicology, conservation biology) [3–5].

In addition, behavior being at the direct interface between organisms and the environment, behavioral changes can drastically affect individual and species interactions [6] through, for instance, alterations in habitat choice, movements, foraging, social and reproductive behavior [7] with expected cascading effects on population dynamics, food webs [8], and ecosystem functions such as primary production, transport of nutrients, and hence biogeochemical cycling [9]. While animal movement and behavioral changes can directly alter ecosystem functioning, they may also prevent rapid population decline through adaptive responses, facilitating genetic adaptation through rapid evolution [1,10], or rather the opposite, resulting in population decline and even extinction when altered in a maladaptive way (*sensu* ecological & evolutionary traps; [1,11,12]). Accordingly, animal movement, migration, and dispersal may indirectly alter ecosystem functioning by affecting eco-evolutionary dynamics [13–15]. For instance, the dispersal rate among metapopulations of baker’s yeast exposed to increasing concentrations of antibiotics has been shown to be an important driver of population adaptation and persistence [16].

Accordingly, improving our understanding of how animals move and behave in various contexts may help to tackle critical conservation aims [17,18], including the need to stabilize or increase the numbers of small or declining populations [19,20], ensure the dispersal of biocontrol agents under various conditions [21,22] and control invasive or pest species [23,24]. For instance, a recent study has shown that combining biopesticides (i.e., essential oils) and biocontrol agents (*Trichogramma evanescens*) to mitigate pest species in agricultural fields may be counter-productive by altering biocontrol agents’ fitness and behavior and hence their efficiency [21].

Overall, what appears as subtle effects on individuals’ behavior may significantly affect ecosystems’ functioning, evolutionary responses, and population persistence, stressing the need to study global changes’ direct and indirect effects on animal behavior. In line with these needs, the increasing use of computer vision approaches, such as automated video-tracking, occurring over the past decade has enabled the collection of high-resolution movement data easily and quickly. In addition, recent methodological and technological innovations, mainly through the implementation of artificial intelligence and machine learning algorithms to video-tracking solutions, have empowered the amount of data collected, allowing, for instance, to track large groups of unmarked individuals simultaneously (e.g., idtracker.ai [26] and TRex [27]). Many open-source tracking solutions now exist and are under active development with great potential for the experimental study of animal behavior (for a recent review, see [28]). However, analyzing and interpreting the data obtained through automated tracking solutions can remain challenging due to their complexity, heterogeneity, and noisiness. In practice, some (possibly substantial) pre-processing is required to make sense of the raw video-tracking data.

To address these challenges and promote the widespread use of video-tracking analyses, we developed MoveR, an R package [29] for importing, filtering, analyzing, and visualizing animal movement and behavior data obtained from common video-tracking solutions such as Ctrax, Idtracker.ai, and TRex. MoveR includes a suite of flexible tools helping to clean data by removing suspected tracking errors/artifacts and computing a range of metrics for quantifying animal movement and behavior (e.g., speed, average neighbor distance). MoveR also allows the computation of the mean and net square displacement values [30,31], which is helpful in inferring population dispersal. In addition, the package includes an original unsupervised learning method relying on density-based clustering [32] for classifying activity states (active vs. inactive) in a two-dimensional space instead of the classic method, generally based on arbitrary thresholds of traveled distance or speed.

To illustrate the utility of MoveR, we present a case study using video-tracking of a group of parasitoid micro-wasps (*Trichogramma sp*.) exposed to a linear temperature ramp increasing from 35 to 45 degrees Celsius and then decreasing back to 35 degrees. We show how MoveR can remove artifactual noise in the data, compute movement and behavior metrics, especially identifying activity states, and thus estimate temperature tolerance thresholds. Our results underline the utility of MoveR in improving the reproducibility and reliability of animal movement and behavior data analyses and in facilitating the deployment of video-tracking techniques in an era of open science.

### 1.2. Related work

With the growing interest in the field of movement and behavioral studies, some proprietary commercial software (e.g., Ethovision [34], LoliTrack - Loligo Systems ApS, Viborg, Denmark), and more recently open source (e.g., AnimalTA [33]), video-tracking solutions have implemented the possibility of running some data processing, computation, and visualization. However, commercial solutions are often expensive, are based on point & click interfaces with limited flexibility, and do not integrate the statistical tools that the constantly evolving R community proposes.

On the other hand, several open-source R packages have been developed. However, they mainly focus on simulating animal movement [35] or analyzing spatial data retrieved from G.P.S. chipsets/collars or A.R.G.O.S. markers by investigating space use and habitat selection by wildlife [36,37], or identifying behavioral states by fitting State–space [38] or hidden Markov models [39,40] (for review see [41]).

To our knowledge, the only R package that preceded MoveR in analyzing animal movement in such a flexible way is trajr [42]. However, while implementing the possibility of computing several interesting metrics (e.g., sinuosity [43,44]) to analyze animal trajectories (i.e., a set of time-specific discrete locations, generally restricted to two spatial dimensions), trajr lacks some useful functionalities that may help both beginners and advanced user, for instance, to import raw tracking data from various tracking software and then filter/clean them into R environment. Accordingly, there is no solution on the R platform to perform pre-processing steps of raw tracking data through detecting and filtering potential tracking artifacts, such as spurious detections or changes in individual identity, which significantly skew results interpretation. In addition, there is no method to handle regions of interest (R.O.I.) or analyze temporal trends across multi-individual tracking sessions, which is crucial for studying behavior in changing environments (see Illustrative examples section below).

For these reasons, MoveR intends to fill the gaps by providing a complete workflow that allows easy import, management, filtering, evaluation, analysis, and visualization of automated video-tracking solutions’ output in a flexible, reproducible, reliable, and open framework as required to push forward the shift toward open science that has been initiated in ecology, evolution and related fields of research [45–47].

## 2. Software description

The MoveR R package is developed within the R environment, an open-source platform [29] continuously evolving, which has been overgrown in the past years through user-contributed packages, especially in ecology [48]. MoveR is fully coded in the base R language, meaning it does not need any compilation. It is highly portable (available for Linux, OS X, and Microsoft Windows) and only relies on few dependencies (Table 1).

**Table 1.**
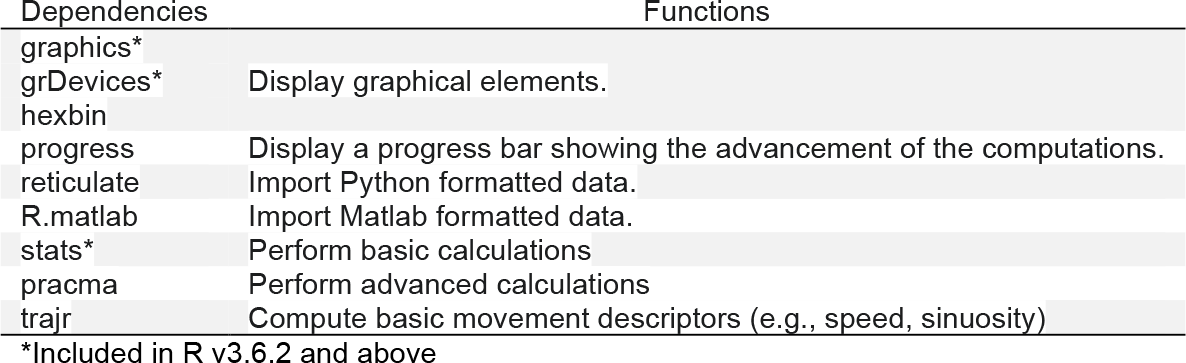
MoveR dependencies.

As the focal object tracked through video-tracking software may be from various taxonomic groups or even be a moving object, we will refer to the term “particle” to specify the focal object/animal tracked hereafter.

### 2.1. Software architecture

MoveR architecture relies on six main functionalities (Figure 1), providing tools (R functions) to (i) import raw data output from various video-tracking software (Figure 1, blue-green box), (ii) visualize pre- and post-processed tracklets (i.e., fragments of the track/trajectory followed by a moving particle), (iii) manage regions of interest (R.O.I), (iv) evaluate the detection rate over the tracking process using particles’ positions manually annotated, (v) clean/filter potential tracking artifact due to the experimental design, the study system (i.e., the focal species) or the tracking algorithm based on user-defined filters (Figure 1, blue boxes). Finally, (vi) the MoveR workflow enables the analysis of particles’ tracklets through the computation of basic or more advanced metrics describing animal movements (Figure 1, purple boxes). For example, advanced functionalities allow the unsupervised classification of activity states, the computation of sociability index or population dispersal (i.e., MSD and Dcoef), the computation of temporal trends, and the identification of particular motifs of behavioral states or changes in areas’ locations. In addition, the resulting output is compatible with various R packages to conduct further statistical tests and modeling classically used in ecology and evolution (e.g., [49]).

**Figure 1.**
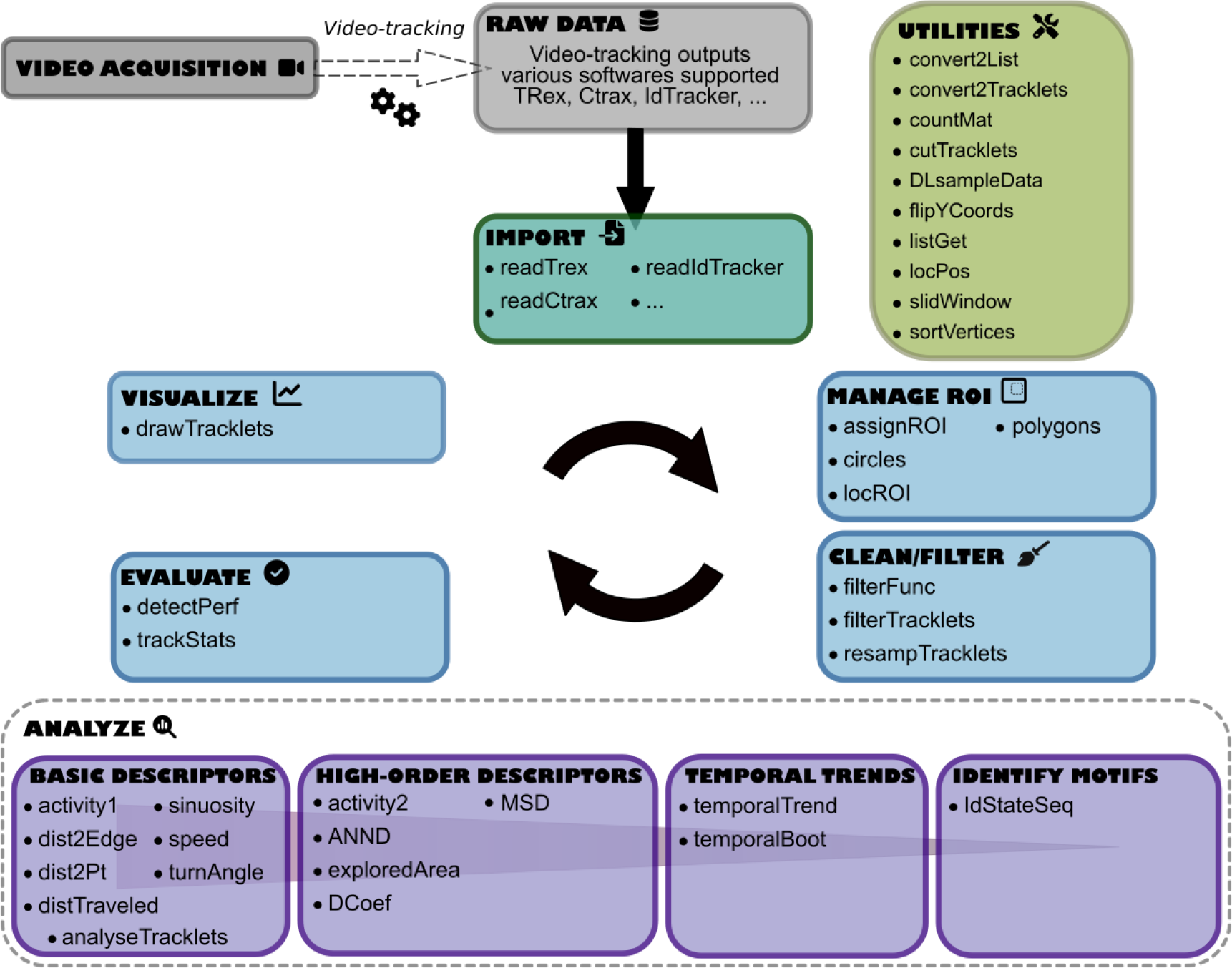
MoveR package workflow and six main functionalities: import, visualize, manage ROI, evaluate, clean/filter, and analyze.

### 2.2. Software functionalities

MoveR provides a highly flexible and unified environment relying on six main functionalities, including explicitly named functions using camelCase programming conventions (see Figure 1 and Table 2).

**Table 2.**
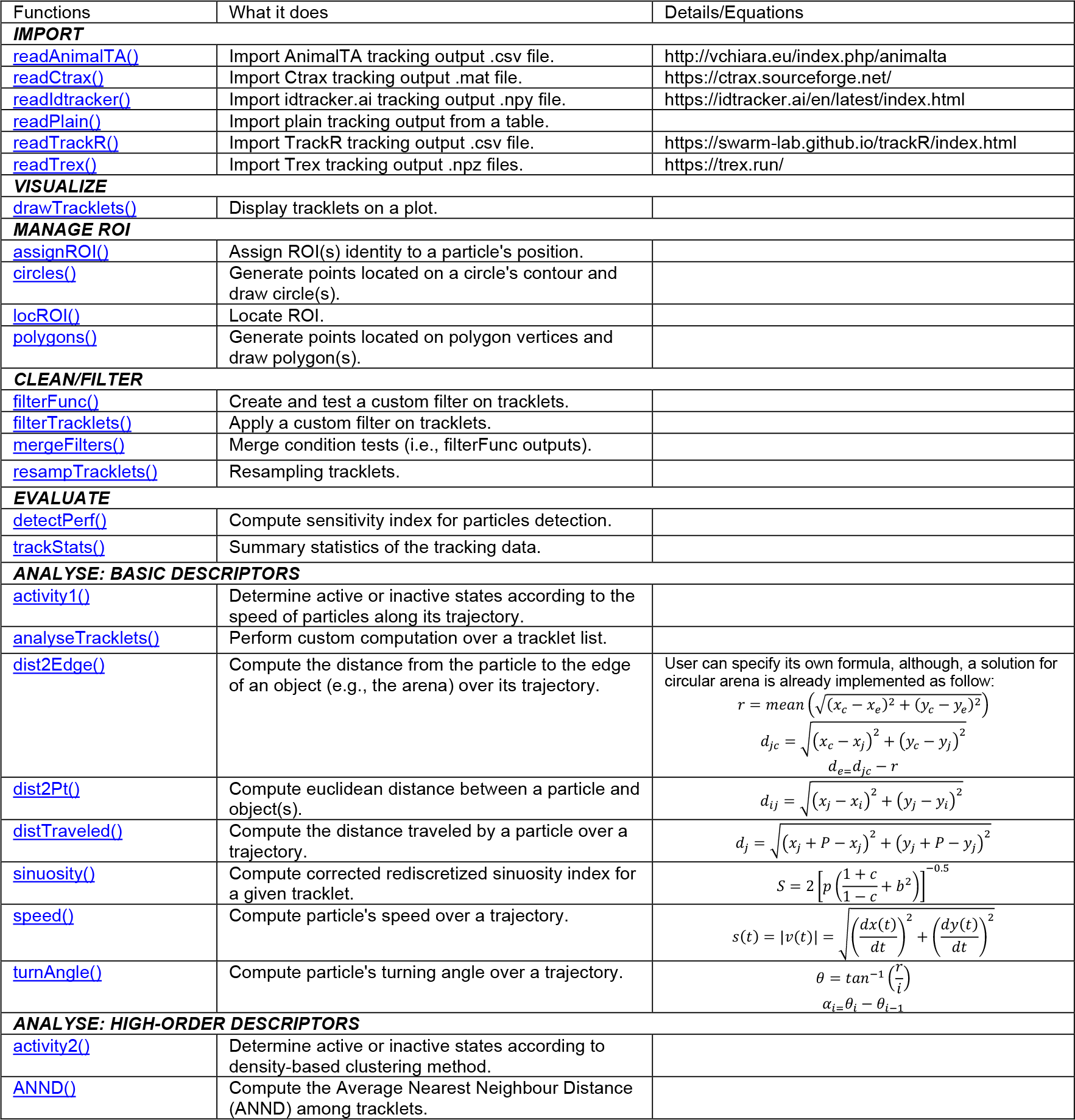
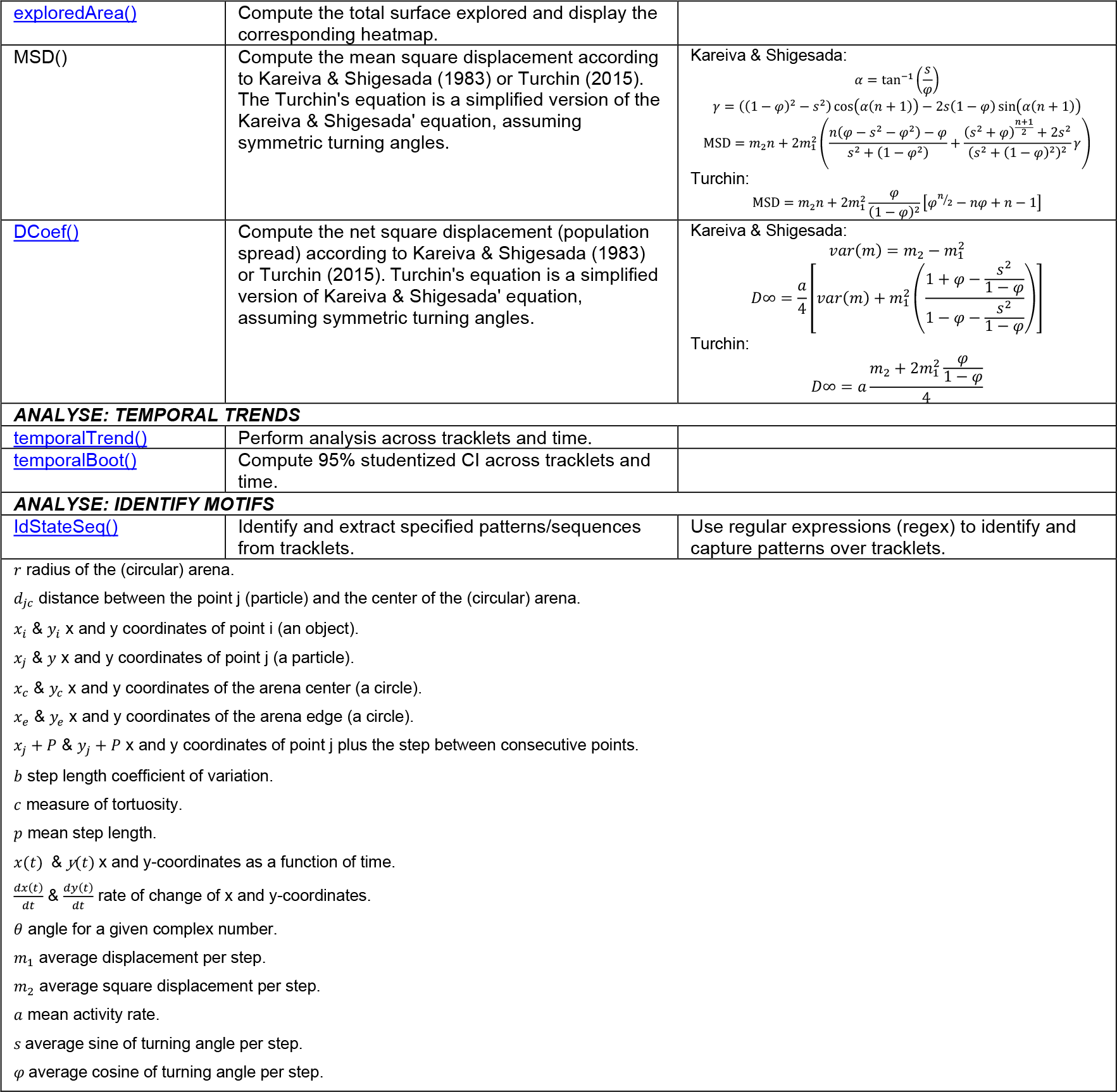
Main functions provided by the MoveR package and quick description of what they help to achieve.

Briefly, **he import** functionality allows raw data integration from software like Ctrax and idtracker.ai using intuitive read functions (e.g., readCtrax).

**The visualize** functionality allows displaying pre- or post-processed tracklets using ‘drawTracklets’ function, which supports various customizations, including adding elements like R.O.I. or arena boundaries.

**The manage R.O.I**. functionality allows importing the location of area edges from a distance matrix generated using an image processing program (e.g., [50]) through the ‘locROI’ function or by specifying it from scratch using ‘circles’ or ‘polygons’ functions. Once R.O.I.s are specified, the ‘assignROI’ function can retrieve the particles’ location among them.

**The clean/filter** functionality allows filtering the data according to user-defined functions using the ‘filterFunc’ function and passed to ‘filterTracklets‘. Indeed, potential tracking artifacts or identity switching may depend on experimental design or a particular system/focal species. A quantitative summary of filtered data is also provided at the end of the filtering process.

**The evaluate** functionality allows computing a quantitative summary of the video and tracking statistics (e.g., video duration, the number of tracklets, and their length and duration) through the call to the ‘trackStats’ function. It also allows to assess video-tracking and filtering process accuracy by comparing manual and automated detections using the ‘detectPerf’ function (i.e., check the amount of true and false detection compared to manual annotation).

**The analyze** functionality is versatile and enables a wide range of calculations, either pre-set (e.g., speed) or user-defined on tracklets, time, or space (e.g., across areas) and behavioral state (e.g., among activity states) using functions like ‘analyseTracklets’ and ‘temporalTrends’ and ‘idStateSeq’. Here, four types of functions can be described:

- Basic descriptors can help to compute classic metrics over tracklets, such as speed and sinuosity.
- High-order descriptors can help to characterize activity states (i.e., active vs. inactive states) using unsupervised learning methods with the ‘activity2’ function according to the density-based clustering method reported in [32] and implemented by [51]. Alternatively, it also makes possible to compute the expected diffusion coefficient, a proxy of population dispersal, assuming a correlated random walk model using the ‘Dcoef’ function according to [30,31].
- The ‘temporalTrend’ function returns the results of a given calculation (weighed by the tracklet length or not) over time by averaging the value for a given time window over each tracklet. Furthermore, computing a studentized 95% confidence interval by bootstrapping over the tracklets is also possible with the ‘temporalBoot’ function.
- The ‘idStateSeq’ function helps to identify and extract arbitrary patterns regarding changes among behavioral states, spatial regions or areas of interest, patch crosses, or any other motifs using a versatile regular-expression syntax.

MoveR offers extensive functionalities addressing various objectives; however, the upcoming example will highlight only common procedures. For the installation procedure, more examples, and tutorials on how to benefit from MoveR’s potential, See the package documentation website built with pkgdown v2.0.7. [52]: https://qpetitjean.github.io/MoveR/.

## 3. Illustrative examples

For this example, we download and import a sample dataset from the MoveR package using the ‘DLsampleData’ and ‘readTrex’ functions (full code is reported in supplementary material).

Briefly, the dataset consists of video recording (1920×1080 pixels) of 24 parasitic micro-wasp individuals (genus *Trichogramma*) placed in a circular arena (diameter: 2.5cm, see [53]) for 110 minutes at 25 fps (see the left panel of Figure 2). Over the exposure duration, the temperature steadily increased from 18°C to 45°C and then decreased back to 18°C (temperature changing rate: 0.5°C per minute). Individuals were then tracked using TRex (v1.1.3) [27]. To reduce space allocation and computing time, we reduced the dataset by removing the movements recorded below 35°C (see the right panel of Figure 2).

**Figure 2.**
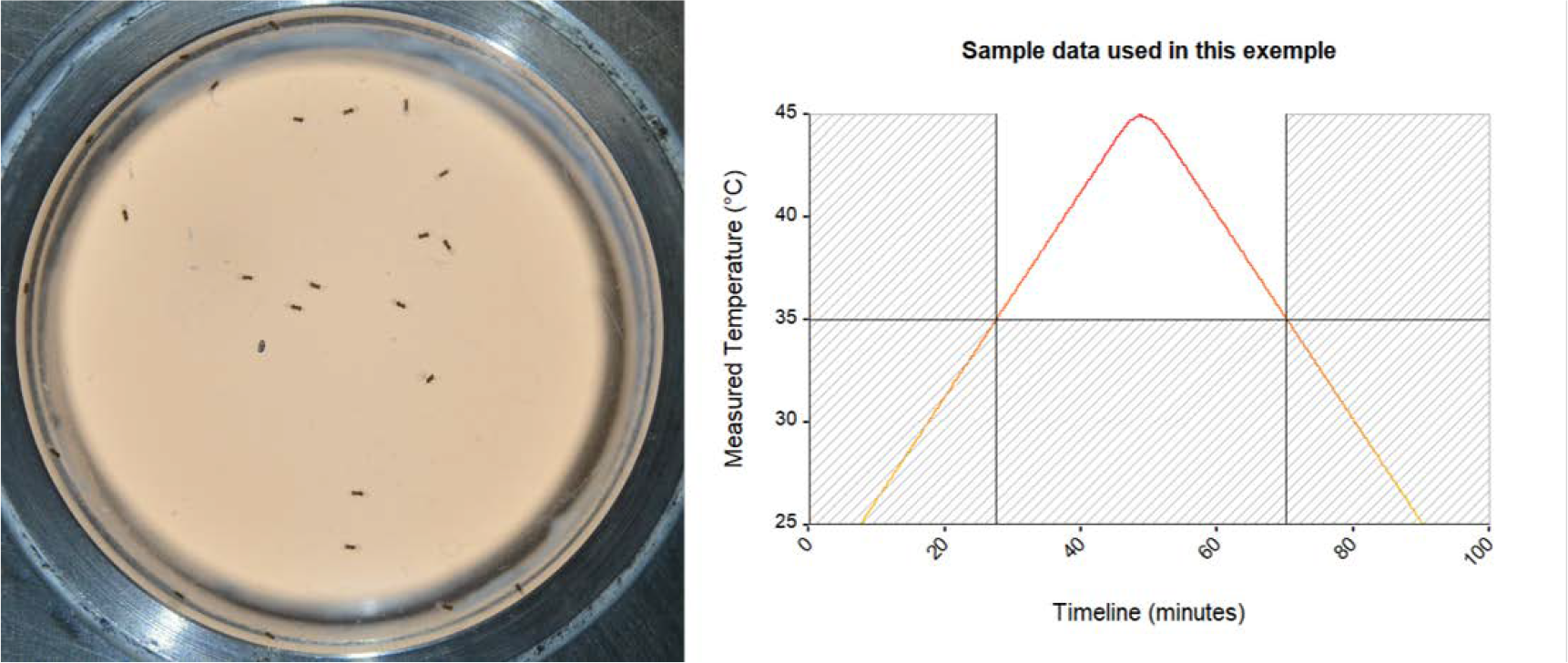
Frames extracted from a video recording of 24 parasitic micro-wasps (genus Trichogramma) placed in a thermostated circular arena (left). Temperature ramp over which the movements of parasitic micro-wasp were recorded. The shaded area corresponds to the part of the data removed from the original dataset (i.e., moments where the temperature was below 35°C), while the white area corresponds to the remaining data used in the following example (right).

Here, we aim to clean the dataset, assign R.O.I. identities to particle locations, compute basic descriptors, and identify activity states (active vs. inactive) on micro-wasps movements to display temporal trends. In other words, we want to identify the moments where the general pattern of micro-wasps movement changes.

After importing (‘readTrex’ function, for code, see supplementary material 2. Importing the data) and cleaning the dataset by removing infinite values (i.e., undetected or lost particles), and spurious detection performed outside the arena (for code, see supplementary material 3. Cleaning the data), we are generating the coordinates of several circular R.O.I. with increasing distance to the center of the arena (for code, see supplementary material 4. Identify the tracklets within an ROI.). Then, using ‘assignROI’, we assign the R.O.I.’s identity to the particles’ position and visualize the tracklets using the ‘drawTracklets’ function (see Figure 3). As the R.O.I.s are circular and the larger ones include smaller ones, a particle detected within the center of the arena (ROI_1 in Figure 3) is also included within the larger R.O.I.s (e.g., particles in ROI_1 are also included in ROI_2, 3 and 4, see Figure 3).

**Figure 3.**
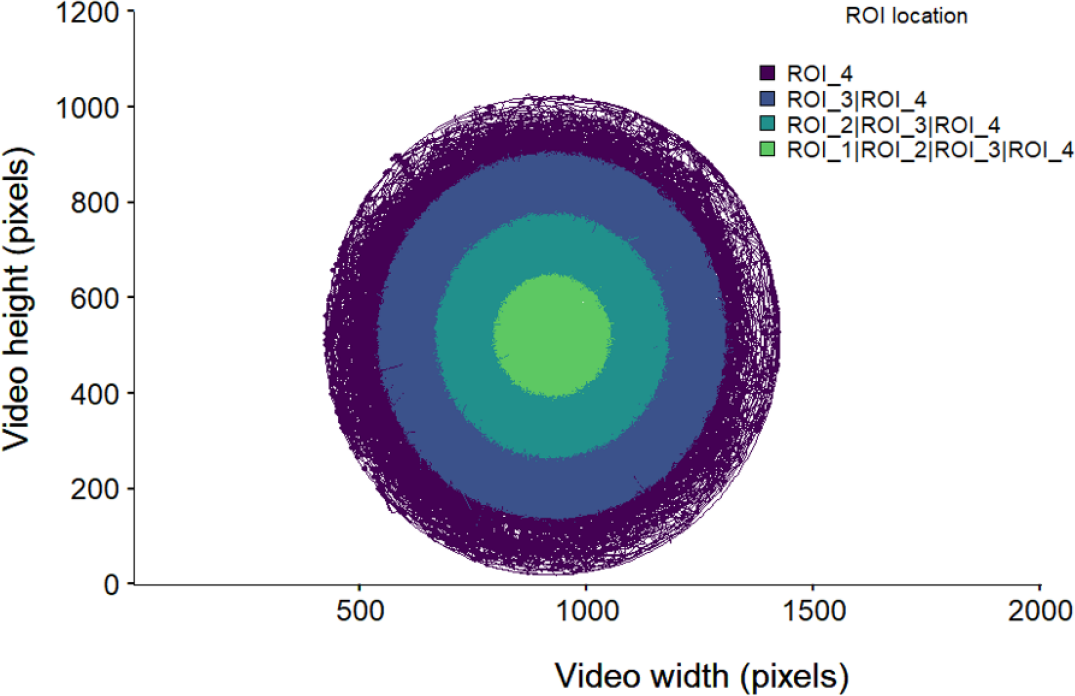
Particle trajectories are colored according to their location among four circular R.O.I.s. Colors indicate the ROI where particles are detected. As larger R.O.I.s include smaller ones, particles located at the arena’s center are considered within the R.O.I. 1, 2, 3, and 4.

Next, we are computing some basic descriptors such as the speed, turning angle, turning angle variance, and sinuosity (using ‘speed’, ‘turnAngle’ and ‘sinuosity’ functions, respectively; for code, see supplementary material 5.1. Compute basic descriptors).

The previous basic descriptors then enable to identify the activity states based in a two-dimensional space (i.e., the speed and the variance of the turning angle; for code, see supplementary material 5.2 Activity states: 2D non-hierarchical clustering) using the ‘activity2’ function. This method uses the density-based clustering algorithm from [23] and implemented by [41] to detect two clusters, assuming inactive states fit a Gaussian distribution. States within the 95% confidence ellipsis or below its major and minor radii are labeled inactive (0), and others as active (1) (See Figure 4).

**Figure 4.**
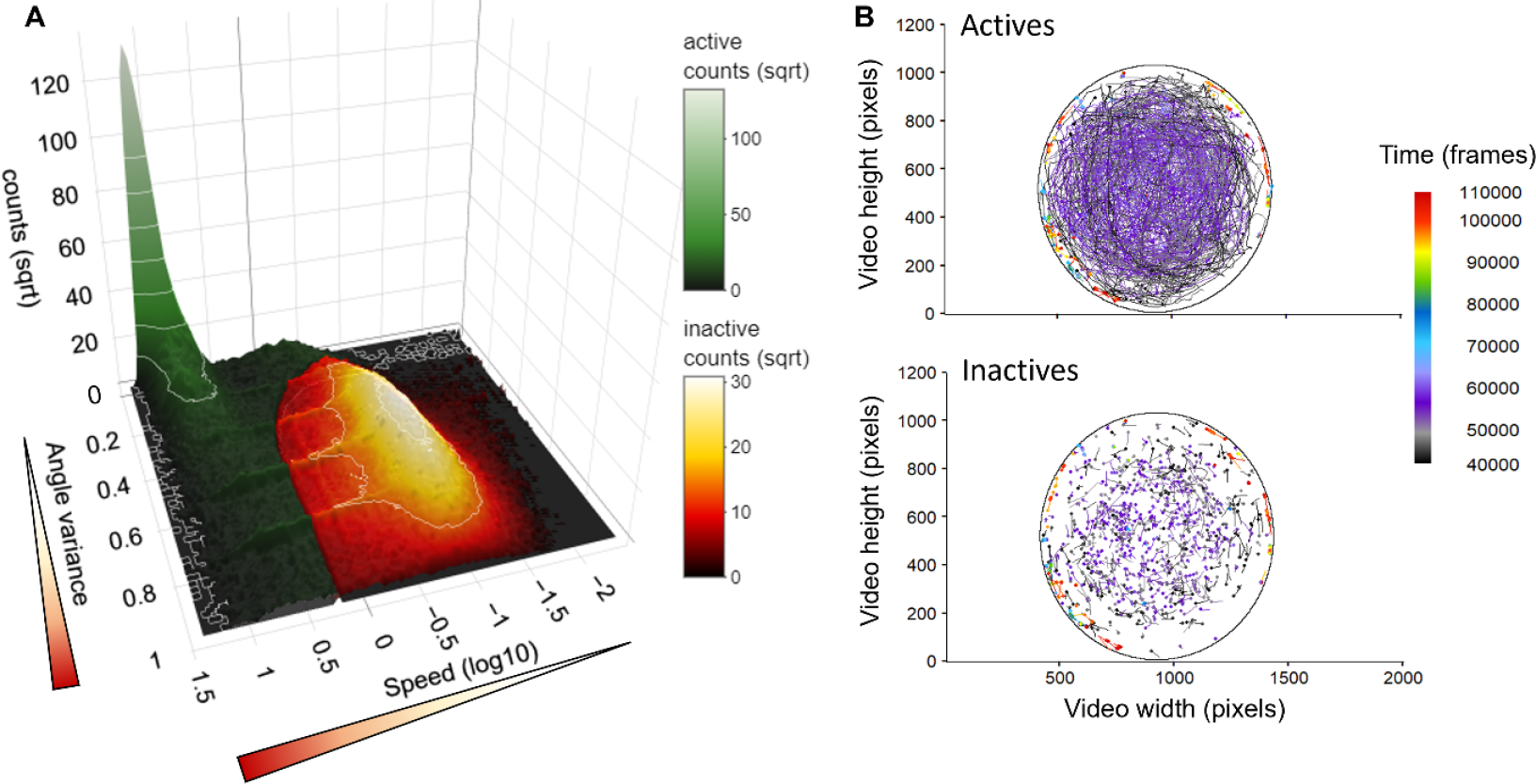
3d density map of the active and inactive states according to the non-hierarchical classification (density-based clustering) performed on smoothed particles speed (log-transformed) and turning angle variance (A) and representation of a sample of active (B, upper panel) and inactive (B lower panel) parts of particles trajectories (i.e., 2000 tracklets). In the 3D density map (A), the active and inactive state distributions are respectively represented as a gradient of dark green to white and dark red to white according to the increasing number of counts. Also, for the representation of the active and inactive parts of a sample of the particles’ trajectories (B), we can observe that the active tracklets are generally longer than the inactive parts.

We then analyze the changes of previous descriptors over time using ‘temporalTrend’ and ‘temporalBoot’ functions, yielding a trend and a studentized 95% confidence interval. (For code, see supplementary material 5.3 Compute and draw temporal trends).

Finally, we can graphically (in this example) retrieve the moment at which movements start to decrease or even stop, as shown by both the evolution of the sinuosity (Figure 5A), the speed (Figure 5B), and the activity rate (Figure 5C) of the individuals over time.

**Figure 5.**
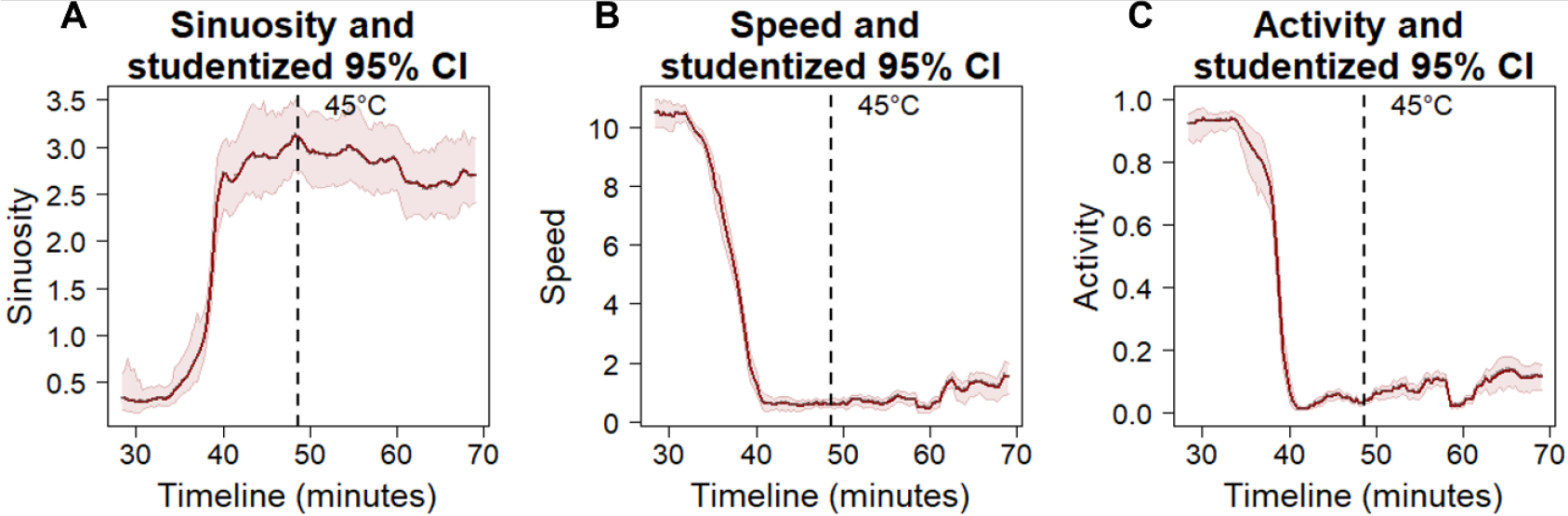
Evolution of the sinuosity (A) speed (B) and activity (C) of Trichogramma individuals (dark red line) and studentized 95% confidence interval (light red envelope) over time (expressed in frame). The dashed line indicates when the maximum temperature has been reached over the ramp: 45°C.

## 4. Impact

The tools developed in the MoveR R package workflow are designed to ease the management and analysis of video-tracking data for beginners and advanced users. By providing an efficient analysis pipeline, MoveR should pave the way for the increasing use of movement and behavioral data to tackle emergent questions encompassing the fields of ecology, evolution, and conservation biology. Here, we provide some examples of applications, but MoveR usages are not limited to these examples and may expand to a broader extent.

More precisely, MoveR can be used to investigate a large array of questions related to, for instance, the variability of responses among species, populations, or individuals by providing the necessary tools for animal video phenotyping. Indeed, while inter-specific variability of responses has been studied extensively, understanding the ecological drivers and effects of intra-specific variability on population dynamics and ecosystem functioning is still an emerging research field that needs further investigations [8,54]. While advances in machine learning algorithms conserving the identity of individuals within a group are growing, MoveR can efficiently allow specifying a reproducible and customized workflow for running computation among several individuals (i.e., tracklets) to, for instance, investigate further questions related to animal personality and syndromes [55–57] under laboratory or mesocosms settings.

In addition, under the context of global changes, the increasing use of synthetic compounds (especially in agricultural fields [58,59]) and the release of micropollutants such as pharmaceuticals [60], endocrine disruptors [61] or nanoparticles [62], and microplastics [63] MoveR represent an efficient tool to assess movements and behavioral alterations caused by such compounds on terrestrial and aquatic fauna. Indeed, the effects of contaminants and, more generally, multiple stressors and their interactive effects on animals’ movement and behavior have largely been overlooked until recently, despite a clear relationship between behavioral alteration and cascading effects on individual and population fitness [4,5,64]. MoveR can hence help ecophysiologists and ecotoxicologists assess the effects of various stress (e.g., xenobiotics, parasitism, temperature changes) in animals by finding relevant and sensitive endpoints related to animal movement and behavior.

Finally, MoveR can be useful for conservation purposes by, for instance, helping to better understand how environmental or social factors affect movements and predict the dispersion of invasive populations (i.e., progression front) [24] or biocontrol agents (i.e., pest-suppression performance) [65] through video-phenotyping methods.

## 5. Conclusions

Collection, analyses, and resulting interpretations of movement and behavioral data overlap the boundaries of several disciplines. Yet, the difficulties encountered in easily and efficiently managing such a large amount of data still prevent us from tackling emerging questions linked to fundamental ecology and evolution and their application to conservation biology. For this reason, giving access to a flexible, reliable, and open framework to the scientific and conservation manager communities is critical. MoveR includes a suite of flexible tools for importing, cleaning, and filtering raw data from several video-tracking softwares and allows to compute basic and more advanced metrics to quantify animal movement and behavior easily within a reproducible environment.

## Acknowledgements

This work was funded by the TripTIC and BIDIME projects (ANR-14-CE18-0002 to JY Rasplus and ANR ECOPHYTO « Maturation »-Grant no. ANR-19-ECOM-0010 to N.R.). We thank Louise VAN OUDENHOVE, Martin BERNET, and Renée LE CLECH for helpful discussions at various stages of this package development.

## Data availability

The data used to illustrate the use of the MoveR package in this article is hosted in the following GitHub repository: https://github.com/qpetitjean/MoveR_SampleData, and can be directly imported into the R environment using the ‘DLsampleData’ function of the MoveR package. The code used to illustrate the use of the MoveR package in this article is included in the supplementary material. However, a more extensive version of this example/tutorial can be found in the “*How to*” section of the package documentation website built with pkgdown v2.0.7. [52]: https://qpetitjean.github.io/MoveR/

## References

[1] B.B.M. Wong, U. Candolin, Behavioral responses to changing environments, Behavioral Ecology. 26 (2015) 665–673. 10.1093/beheco/aru183.

[2] T. Rahman, U. Candolin, Linking animal behavior to ecosystem change in disturbed environments, Front. Ecol. Evol. 10 (2022). 10.3389/fevo.2022.893453.

[3] E.E. Little, S.E. Finger, Swimming behavior as an indicator of sublethal toxicity in fish, Environmental Toxicology and Chemistry. 9 (1990) 13–19. 10.1002/etc.5620090103.

[4] E.K. Peterson, D.B. Buchwalter, J.L. Kerby, M.K. LeFauve, C.W. Varian-Ramos, J.P. Swaddle, Integrative behavioral ecotoxicology: bringing together fields to establish new insight to behavioral ecology, toxicology, and conservation, Curr Zool. 63 (2017) 185–194. 10.1093/cz/zox010.

[5] M.G. Bertram, J.M. Martin, E.S. McCallum, L.A. Alton, J.A. Brand, B.W. Brooks, D. Cerveny, J. Fick, A.T. Ford, G. Hellström, M. Michelangeli, S. Nakagawa, G. Polverino, M. Saaristo, A. Sih, H. Tan, C.R. Tyler, B.B.M. Wong, T. Brodin, Frontiers in quantifying wildlife behavioural responses to chemical pollution, Biological Reviews. n/a (2022). 10.1111/brv.12844.

[6] T.J. Bartley, K.S. McCann, C. Bieg, K. Cazelles, M. Granados, M.M. Guzzo, A.S. MacDougall, T.D. Tunney, B.C. McMeans, Food web rewiring in a changing world, Nat Ecol Evol. 3 (2019) 345–354. 10.1038/s41559-018-0772-3.

[7] U. Tuomainen, U. Candolin, Behavioural responses to human-induced environmental change, Biol Rev Camb Philos Soc. 86 (2011) 640–657. 10.1111/j.1469-185X.2010.00164.x.

[8] N.P. Moran, B.B.M. Wong, R.M. Thompson, Weaving animal temperament into food webs: implications for biodiversity, Oikos. 126 (2017) 917–930. 10.1111/oik.03642.

[9] E.S. Bakker, K.A. Wood, J.F. Pagès, G.F. (Ciska) Veen, M.J.A. Christianen, L. Santamaría, B.A. Nolet, S. Hilt, Herbivory on freshwater and marine macrophytes: A review and perspective, Aquatic Botany. 135 (2016) 18–36. 10.1016/j.aquabot.2016.04.008.

[10] V. Calcagno, M. Dubosclard, C. de Mazancourt, Rapid Exploiter-Victim Coevolution: The Race Is Not Always to the Swift., The American Naturalist. 176 (2010) 198–211. 10.1086/653665.

[11] B.A. Robertson, J.S. Rehage, A. Sih, Ecological novelty and the emergence of evolutionary traps, Trends in Ecology & Evolution. 28 (2013) 552–560. 10.1016/j.tree.2013.04.004.

[12] M.A. Schlaepfer, M.C. Runge, P.W. Sherman, Ecological and evolutionary traps, Trends in Ecology & Evolution. 17 (2002) 474–480. 10.1016/S0169-5347(02)02580-6.

[13] G. Bell, Evolutionary Rescue, Annual Review of Ecology, Evolution, and Systematics. 48 (2017) 605– 627. 10.1146/annurev-ecolsys-110316-023011.

[14] F. Aubree, B. Lac, L. Mailleret, V. Calcagno, Migration pulsedness alters patterns of allele fixation and local adaptation in a mainland-island model, Evolution. 77 (2023) 718–730. 10.1093/evolut/qpac067.

[15] F. Aubree, P. David, P. Jarne, M. Loreau, N. Mouquet, V. Calcagno, How community adaptation affects biodiversity–ecosystem functioning relationships, Ecology Letters. 23 (2020) 1263–1275. 10.1111/ele.13530.

[16] G. Bell, A. Gonzalez, Adaptation and Evolutionary Rescue in Metapopulations Experiencing Environmental Deterioration, Science. 332 (2011) 1327–1330. 10.1126/science.1203105.

[17] A.L. Greggor, D.T. Blumstein, B.B.M. Wong, O. Berger-Tal, Using animal behavior in conservation management: a series of systematic reviews and maps, Environmental Evidence. 8 (2019) 23. 10.1186/s13750-019-0164-4.

[18] K.C. Fraser, K.T.A. Davies, C.M. Davy, A.T. Ford, D.T.T. Flockhart, E.G. Martins, Tracking the Conservation Promise of Movement Ecology, Frontiers in Ecology and Evolution. 6 (2018). https://www.frontiersin.org/articles/10.3389/fevo.2018.00150 (accessed June 2, 2023).

[19] J. Clobert, J.-F. Le Galliard, J. Cote, S. Meylan, M. Massot, Informed dispersal, heterogeneity in animal dispersal syndromes and the dynamics of spatially structured populations, Ecology Letters. 12 (2009) 197–209. 10.1111/j.1461-0248.2008.01267.x.

[20] J. Cote, J. Clobert, Social personalities influence natal dispersal in a lizard, Proceedings of the Royal Society B: Biological Sciences. 274 (2006) 383–390. 10.1098/rspb.2006.3734.

[21] L. van Oudenhove, A. Cazier, M. Fillaud, A.-V. Lavoir, H. Fatnassi, G. Perez, V. Calcagno, Non-target effects of ten essential oils on the egg parasitoid Trichogramma evanescens, Peer Community Journal. 3 (2023). 10.24072/pcjournal.212.

[22] M. Cointe, V. Burte, G. Perez, L. Mailleret, V. Calcagno, A double-spiral maze and hi-resolution tracking pipeline to study dispersal by groups of minute insects, Sci Rep. 13 (2023) 5200. 10.1038/s41598-023-31630-8.

[23] R.A. Duckworth, A.V. Badyaev, Coupling of dispersal and aggression facilitates the rapid range expansion of a passerine bird, Proceedings of the National Academy of Sciences. 104 (2007) 15017–15022. 10.1073/pnas.0706174104.

[24] J. Cote, S. Fogarty, K. Weinersmith, T. Brodin, A. Sih, Personality traits and dispersal tendency in the invasive mosquitofish (Gambusia affinis), Proceedings of the Royal Society B: Biological Sciences. 277 (2010) 1571–1579. 10.1098/rspb.2009.2128.

[25] O. Berger-Tal, T. Polak, A. Oron, Y. Lubin, B.P. Kotler, D. Saltz, Integrating animal behavior and conservation biology: a conceptual framework, Behavioral Ecology. 22 (2011) 236–239. 10.1093/beheco/arq224.

[26] F. Romero-Ferrero, M.G. Bergomi, R.C. Hinz, F.J.H. Heras, G.G. de Polavieja, idtracker.ai: tracking all individuals in small or large collectives of unmarked animals, Nat Methods. 16 (2019) 179–182. 10.1038/s41592-018-0295-5.

[27] T. Walter, I.D. Couzin, TRex, a fast multi-animal tracking system with markerless identification, and 2D estimation of posture and visual fields, eLife. 10 (2021) e64000. 10.7554/eLife.64000.

[28] V. Panadeiro, A. Rodriguez, J. Henry, D. Wlodkowic, M. Andersson, A review of 28 free animal-tracking software applications: current features and limitations, Lab Anim. (2021) 1–9. 10.1038/s41684-021-00811-1.

[29] R Core Team, R: A language and environment for statistical computing. R Foundation for Statistical Computing, Vienna, Austria., (2020). https://www.R-project.org/.

[30] P. Turchin, Quantitative Analysis of Movement: Measuring and Modeling Population Redistribution in Animals and Plants, Beresta Books, Storrs, Connecticut, 2015.

[31] P.M. Kareiva, N. Shigesada, Analyzing insect movement as a correlated random walk, Oecologia. 56 (1983) 234–238. 10.1007/BF00379695.

[32] M. Ester, H.-P. Kriegel, J. Sander, X. Xu, A density-based algorithm for discovering clusters in large spatial databases with noise, in: Proceedings of the Second International Conference on Knowledge Discovery and Data Mining, AAAI Press, Portland, Oregon, 1996: pp. 226–231.

[33] V. Chiara, S.-Y. Kim, AnimalTA: A highly flexible and easy-to-use program for tracking and analysing animal movement in different environments, Methods in Ecology and Evolution. 14 (2023) 1699– 1707. 10.1111/2041-210X.14115.

[34] L.P.J.J. Noldus, A.J. Spink, R.A.J. Tegelenbosch, EthoVision: A versatile video tracking system for automation of behavioral experiments, Behavior Research Methods, Instruments, & Computers. 33 (2001) 398–414. 10.3758/BF03195394.

[35] B.M. Marshall, A.B. Duthie, abmAnimalMovement: An R package for simulating animal movement using an agent-based model, (2022). 10.12688/f1000research.124810.1.

[36] C. Calenge, The package “adehabitat” for the R software: A tool for the analysis of space and habitat use by animals, Ecological Modelling. 197 (2006) 516–519. 10.1016/j.ecolmodel.2006.03.017.

[37] C. Calenge, S. Dray, M. Royer-Carenzi, The concept of animals’ trajectories from a data analysis perspective, Ecological Informatics. 4 (2009) 34–41. 10.1016/j.ecoinf.2008.10.002.

[38] I.D. Jonsen, W.J. Grecian, L. Phillips, G. Carroll, C. McMahon, R.G. Harcourt, M.A. Hindell, T.A. Patterson, aniMotum, an R package for animal movement data: Rapid quality control, behavioural estimation and simulation, Methods in Ecology and Evolution. 14 (2023) 806–816. 10.1111/2041-210X.14060.

[39] T. Michelot, R. Langrock, T.A. Patterson, moveHMM: an R package for the statistical modelling of animal movement data using hidden Markov models, Methods in Ecology and Evolution. 7 (2016) 1308–1315. 10.1111/2041-210X.12578.

[40] B.T. McClintock, T. Michelot, momentuHMM: R package for generalized hidden Markov models of animal movement, Methods in Ecology and Evolution. 9 (2018) 1518–1530. 10.1111/2041-210X.12995.

[41] R. Joo, M.E. Boone, T.A. Clay, S.C. Patrick, S. Clusella-Trullas, M. Basille, Navigating through the r packages for movement, Journal of Animal Ecology. 89 (2020) 248–267. 10.1111/1365-2656.13116.

[42] D.J. McLean, M.A. Skowron Volponi, trajr: An R package for characterisation of animal trajectories, Ethology. 124 (2018) 440–448. 10.1111/eth.12739.

[43] S. Benhamou, How to reliably estimate the tortuosity of an animal’s path:, Journal of Theoretical Biology. 229 (2004) 209–220. 10.1016/j.jtbi.2004.03.016.

[44] P. Bovet, S. Benhamou, Spatial analysis of animals’ movements using a correlated random walk model, Journal of Theoretical Biology. 131 (1988) 419–433. 10.1016/S0022-5193(88)80038-9.

[45] E.M. Wolkovich, J. Regetz, M.I. O’Connor, Advances in global change research require open science by individual researchers, Global Change Biology. 18 (2012) 2102–2110. 10.1111/j.1365-2486.2012.02693.x.

[46] R.E. O’Dea, T.H. Parker, Y.E. Chee, A. Culina, S.M. Drobniak, D.H. Duncan, F. Fidler, E. Gould, M. Ihle, C.D. Kelly, M. Lagisz, D.G. Roche, A. Sánchez-Tójar, D.P. Wilkinson, B.C. Wintle, S. Nakagawa, Towards open, reliable, and transparent ecology and evolutionary biology, BMC Biology. 19 (2021) 68. 10.1186/s12915-021-01006-3.

[47] S.M. Powers, S.E. Hampton, Open science, reproducibility, and transparency in ecology, Ecological Applications. 29 (2019) e01822. 10.1002/eap.1822.

[48] J. Lai, C.J. Lortie, R.A. Muenchen, J. Yang, K. Ma, Evaluating the popularity of R in ecology, Ecosphere. 10 (2019) e02567. 10.1002/ecs2.2567.

[49] D. Bates, M. Maechler, B. Bolker, S. Walker, Fitting Linear Mixed-Effects Models Using lme4, Journal of Statistical Software. 67(1) (2015) 1–48. 10.18637/jss.v067.i01.

[50] C.A. Schneider, W.S. Rasband, K.W. Eliceiri, NIH Image to ImageJ: 25 years of image analysis, Nat Methods. 9 (2012) 671–675. 10.1038/nmeth.2089.

[51] C. Hennig, fpc: Flexible Procedures for Clustering, (2023). https://cran.r-project.org/web/packages/fpc/index.html (accessed June 2, 2023).

[52] H. Wickham, J. Hesselberth, M. Salmon, pkgdown: Make Static HTML Documentation for a Package, (2022). https://pkgdown.r-lib.org,https://github.com/r-lib/pkgdown.

[53] M. Ion Scotta, L. Margris, N. Sellier, S. Warot, F. Gatti, F. Siccardi, P. Gibert, E. Vercken, N. Ris, Genetic Variability, Population Differentiation, and Correlations for Thermal Tolerance Indices in the Minute Wasp, Trichogramma cacoeciae, Insects. 12 (2021) 1013. 10.3390/insects12111013.

[54] A. Raffard, J. Cucherousset, J.G. Prunier, G. Loot, F. Santoul, S. Blanchet, Variability of functional traits and their syndromes in a freshwater fish species (Phoxinus phoxinus): The role of adaptive and nonadaptive processes, Ecol Evol. 9 (2019) 2833–2846. 10.1002/ece3.4961.

[55] S.S. Killen, S. Marras, N.B. Metcalfe, D.J. McKenzie, P. Domenici, Environmental stressors alter relationships between physiology and behaviour, Trends in Ecology & Evolution. 28 (2013) 651– 658. 10.1016/j.tree.2013.05.005.

[56] D. Réale, S.M. Reader, D. Sol, P.T. McDougall, N.J. Dingemanse, Integrating animal temperament within ecology and evolution, Biological Reviews. 82 (2007) 291–318. 10.1111/j.1469-185X.2007.00010.x.

[57] D. Réale, D. Garant, M.M. Humphries, P. Bergeron, V. Careau, P.-O. Montiglio, Personality and the emergence of the pace-of-life syndrome concept at the population level, Philosophical Transactions of the Royal Society B: Biological Sciences. 365 (2010) 4051–4063. 10.1098/rstb.2010.0208.

[58] E.S. Bernhardt, E.J. Rosi, M.O. Gessner, Synthetic chemicals as agents of global change, Frontiers in Ecology and the Environment. 15 (2017) 84–90. 10.1002/fee.1450.

[59] UNEP, Global chemicals outlook, Geneva, Switzerland: United Nations Environment Programme. (2019). https://www.unep.org/explore-topics/chemicals-waste/what-we-do/policy-and-governance/global-chemicals-outlook.

[60] T. Brodin, S. Piovano, J. Fick, J. Klaminder, M. Heynen, M. Jonsson, Ecological effects of pharmaceuticals in aquatic systems—impacts through behavioural alterations, Philos Trans R Soc Lond B Biol Sci. 369 (2014). 10.1098/rstb.2013.0580.

[61] C.D. Metcalfe, S. Bayen, M. Desrosiers, G. Muñoz, S. Sauvé, V. Yargeau, An introduction to the sources, fate, occurrence and effects of endocrine disrupting chemicals released into the environment, Environmental Research. 207 (2022) 112658. 10.1016/j.envres.2021.112658.

[62] M. Bundschuh, F. Seitz, R.R. Rosenfeldt, R. Schulz, Effects of nanoparticles in fresh waters: risks, mechanisms and interactions, Freshwater Biology. 61 (2016) 2185–2196. 10.1111/fwb.12701.

[63] R. Talbot, H. Chang, Microplastics in freshwater: A global review of factors affecting spatial and temporal variations, Environmental Pollution. 292 (2022) 118393. 10.1016/j.envpol.2021.118393.

[64] M. Ågerstrand, K. Arnold, S. Balshine, T. Brodin, B. W. Brooks, G. Maack, E. S. McCallum, G. Pyle, M. Saaristo, A. T. Ford, Emerging investigator series: use of behavioural endpoints in the regulation of chemicals, Environmental Science: Processes & Impacts. 22 (2020) 49–65. 10.1039/C9EM00463G.

[65] V. Burte, M. Cointe, G. Perez, L. Mailleret, V. Calcagno, When complex movement yields simple dispersal: behavioural heterogeneity, spatial spread and parasitism in groups of micro-wasps, Mov Ecol. 11 (2023) 13. 10.1186/s40462-023-00371-8.

