## Supplementary material for "MoveR: an R package for easy processing and analysis of animal video-tracking data": Supp_Mat_MoveR.html

#### 2023-11-08

- 1 Installing MoveR
- 2 Importing the data
  - 2.1 Download the sample dataset
  - 2.2 Import the sample data into R env.
  - 2.3 Retrieve some data
  - 2.4 Retrieve the frame rate of the video (fps)
  - 2.5 Convert the data to a list of tracklets
- 3 Cleaning the data
  - 3.1 Remove infinite values
    - 3.1.1 Specify the filter to detect infinite values on “x.pos”
    - 3.1.2 Specify the filter to detect infinite values on “y.pos”
    - 3.1.3 Merge the previously specified filters
    - 3.1.4 Filter infinite values according to the previously specified filters
    - 3.1.5 Display the summary of the filtering process
  - 3.2 Remove individuals detected outside the arena
    - 3.2.1 Import the location of the arena edge
    - 3.2.2 Draw the tracklets and add the contour of the arena
    - 3.2.3 Assign The ROI (here the arena) to the individuals’ positions
    - 3.2.4 Specify the filter to detect individuals outside the arena
    - 3.2.5 Filter individuals outside the arena according to the previously specified filters
- 4 Identify the tracklets within an ROI
  - 4.1 compute the radius of the arena in pixels
  - 4.2 Generate ROIs coordinates
  - 4.3 Assign ROIs’ identity to individual’s position
  - 4.4 Display colored tracklet according to the area where they are located
- 5 Compute some metrics
  - 5.1 Compute basic descriptors
  - 5.2 Activity states: 2D non-hierarchical clustering
    - 5.2.1 Run the classification
    - 5.2.2 Display the resulting clusters
  - 5.3 Compute and draw temporal trends
    - 5.3.1 Temporal trends
    - 5.3.2 Studentized 95% confidence interval (CI)
    - 5.3.3 Plot the trends & studentized 95% CIs

### 1 Installing MoveR

You can install the development version of MoveR from GitHub with:

```
# install.packages("remotes")
remotes::install_github("qpetitjean/MoveR")
```

Then you can attach the package MoveR:

```
library("MoveR")
```

### 2 Importing the data

#### 2.1 Download the sample dataset

*NB: here we are downloading the second sample dataset made available to depict the use of the MoveR package.*

```
Path2Data <- MoveR::DLsampleData(dataSet = 2, 
                                 tracker = "TRex")
Path2Data
```

```
## [1] "C:\\Users\\quent\\AppData\\Local\\Temp\\RtmpyK4pS0\\MoveR_SampleData-main\\sample_2\\TRexOutput"                                                          
## [2] "C:\\Users\\quent\\AppData\\Local\\Temp\\RtmpyK4pS0\\MoveR_SampleData-main\\sample_2\\ReferenceData\\DistMatrixFromArenaEdge_2021-03-01-ISA17150-chaud.txt"
## [3] "C:\\Users\\quent\\AppData\\Local\\Temp\\RtmpyK4pS0\\MoveR_SampleData-main\\sample_2\\cleaned_2021-03-01-ISA17150-chaud.csv.gz"
```

#### 2.2 Import the sample data into R env.

The import functionality allows importing raw data from video-tracking software such as TRex, Ctrax, and idtracker.ai in the R environment using intuitive `read` functions (e.g., readTrex, readCtrax, readIdTracker).

The data are imported as a list of 9 vectors classically used for further computations using the MoveR package.
\* maj.ax, corresponding to the length of the major axis (i.e., the midline) for a particle over a timeline (i.e., the ellipse’s length).

- angle, corresponding to the particle’s absolute angle expressed in radians (i.e., the particle’s orientation according to the X-axis).
- min.ax, corresponding to the minor axis length for a particle over a timeline (i.e., the ellipse’s width).
- x.pos, corresponding to the x position of the particle’s centroid.
- y.pos, corresponding to the y position of the particle’s centroid.
- identity, corresponding to the particle’s identity given by the tracking software.
- frame, corresponding to the video frame number at which the detection has been performed.
- ntargets, corresponding to the number of particles tracked over the timeline.
- timestamps, corresponding to the elapsed time over each timeline unit in seconds.

*NB: If the raw data are not fully reported within these nine vectors, the “read” functions include a `rawDat` argument which returns a secondary list containing all measures performed by the selected tracking software.*

```
TRexDat <- MoveR::readTrex(Path2Data[[1]],
                           rawDat = T)
str(TRexDat)
```

```
## List of 2
##  $ Data_Trex    :List of 9
##   ..$ maj.ax    : num [1:4716398] 23.8 Inf Inf Inf Inf ...
##   ..$ angle     : num [1:4716398] 1.33 Inf Inf Inf Inf ...
##   ..$ min.ax    : num [1:4716398] NA NA NA NA NA NA NA NA NA NA ...
##   ..$ x.pos     : num [1:4716398] 1372 Inf Inf Inf Inf ...
##   ..$ y.pos     : num [1:4716398] 547 Inf Inf Inf Inf ...
##   ..$ identity  : num [1:4716398] 434 434 434 434 434 434 434 434 434 434 ...
##   ..$ frame     : num [1:4716398] 93065 93066 93067 93068 93069 ...
##   ..$ ntargets  : num [1:64076] 43 25 33 30 31 26 28 28 30 31 ...
##   ..$ timestamps: num [1:64076] 3723 3723 3723 3723 3723 ...
##  $ Data_Trex_Raw:List of 40
##   ..$ BORDER_DISTANCE#pcentroid: num [1:4716398] 535 Inf Inf Inf Inf ...
##   ..$ ACCELERATION#wcentroid   : num [1:4716398] 0 Inf Inf Inf Inf ...
##   ..$ ACCELERATION#pcentroid   : num [1:4716398] 0 Inf Inf Inf Inf ...
##   ..$ SPEED.smooth#wcentroid   : num [1:4716398] 0 Inf Inf Inf Inf ...
##   ..$ Measured_Temp_Deg_C      : num [1:4716398] 39.1 39.1 39.1 39.1 39.1 39.1 39.1 39.1 39.1 39.1 ...
##   ..$ ANGULAR_V#centroid       : num [1:4716398] 0 Inf Inf Inf Inf ...
##   ..$ ANGULAR_A#centroid       : num [1:4716398] 0 Inf Inf Inf Inf ...
##   ..$ normalized_midline       : num [1:4716398] Inf Inf Inf Inf Inf ...
##   ..$ SPEED#wcentroid          : num [1:4716398] 0 Inf Inf Inf Inf ...
##   ..$ SPEED#pcentroid          : num [1:4716398] 0 Inf Inf Inf Inf ...
##   ..$ MIDLINE_OFFSET           : num [1:4716398] -0.0342 Inf Inf Inf Inf ...
##   ..$ midline_length           : num [1:4716398] 23.8 Inf Inf Inf Inf ...
##   ..$ segment_length           : num [1:4716398] 0.991 Inf Inf Inf Inf ...
##   ..$ runTimelinef             : num [1:4716398] 93065 93066 93067 93068 93069 ...
##   ..$ runTimelineS             : num [1:4716398] 3723 3723 3723 3723 3723 ...
##   ..$ RtimeSRound              : num [1:4716398] 46263 46263 46263 46263 46263 ...
##   ..$ X#wcentroid              : num [1:4716398] 1372 Inf Inf Inf Inf ...
##   ..$ Y#wcentroid              : num [1:4716398] 547 Inf Inf Inf Inf ...
##   ..$ Video_Numb               : num [1:4716398] 7 7 7 7 7 7 7 7 7 7 ...
##   ..$ num_pixels               : num [1:4716398] 71 Inf Inf Inf Inf ...
##   ..$ RtimeRound               : num [1:4716398] 2 2 2 2 2 2 2 2 2 2 ...
##   ..$ midline_x                : num [1:4716398] 1375 Inf Inf Inf Inf ...
##   ..$ midline_y                : num [1:4716398] 557 Inf Inf Inf Inf ...
##   ..$ timestamp                : num [1:4716398] 3.72e+09 3.72e+09 3.72e+09 3.72e+09 3.72e+09 ...
##   ..$ videotime                : num [1:4716398] 2.6 2.64 2.68 2.72 2.76 ...
##   ..$ Timeline                 : chr [1:4716398] "12H 51M 2.6S" "12H 51M 2.64S" "12H 51M 2.68S" "12H 51M 2.72S" ...
##   ..$ missing                  : num [1:4716398] 0 1 1 1 1 1 1 1 1 1 ...
##   ..$ IdVidId                  : chr [1:4716398] "434_7" "434_7" "434_7" "434_7" ...
##   ..$ RtimeS                   : num [1:4716398] 46263 46263 46263 46263 46263 ...
##   ..$ frame                    : num [1:4716398] 93065 93066 93067 93068 93069 ...
##   ..$ ANGLE                    : num [1:4716398] 1.33 Inf Inf Inf Inf ...
##   ..$ SPEED                    : num [1:4716398] 0 Inf Inf Inf Inf ...
##   ..$ Rtime                    : num [1:4716398] 2.6 2.64 2.68 2.72 2.76 2.8 2.84 2.88 2.92 2.96 ...
##   ..$ time                     : num [1:4716398] 3723 3723 3723 3723 3723 ...
##   ..$ AY                       : num [1:4716398] 0 Inf Inf Inf Inf ...
##   ..$ AX                       : num [1:4716398] 0 Inf Inf Inf Inf ...
##   ..$ VX                       : num [1:4716398] 0 Inf Inf Inf Inf ...
##   ..$ VY                       : num [1:4716398] 0 Inf Inf Inf Inf ...
##   ..$ X                        : num [1:4716398] 1374 Inf Inf Inf Inf ...
##   ..$ Y                        : num [1:4716398] 554 Inf Inf Inf Inf ...
```

#### 2.3 Retrieve some data

*NB: here we are retrieving some data specifically related to the dataset and that are stored in the Data\_Trex\_Raw sublist: the timeline of the experiment expressed in frames (runTimelinef) and the associated temperature ramp (Measured\_Temp\_Deg\_C).*

```
TRexDat <- c(TRexDat[[1]],
             TRexDat[[2]]["runTimelinef"],
             TRexDat[[2]]["Measured_Temp_Deg_C"])
```

#### 2.4 Retrieve the frame rate of the video (fps)

```
frameRate <- max(TRexDat[["runTimelinef"]], na.rm=T) / max(TRexDat[["timestamps"]], na.rm=T)
```

#### 2.5 Convert the data to a list of tracklets

*NB: at this step also remove ntargets and timestamps because they are not useful for the following steps and their length is lower than other variables.*

```
trackDat <- MoveR::convert2Tracklets(TRexDat[-which(names(TRexDat) == "ntargets" | names(TRexDat) == "timestamps")], by = "identity")
str(trackDat[1:2]) # display only the first two tracklets
```

```
## List of 2
##  $ 434:'data.frame': 12074 obs. of  9 variables:
##   ..$ maj.ax             : num [1:12074] 23.8 Inf Inf Inf Inf ...
##   ..$ angle              : num [1:12074] 1.33 Inf Inf Inf Inf ...
##   ..$ min.ax             : num [1:12074] NA NA NA NA NA NA NA NA NA NA ...
##   ..$ x.pos              : num [1:12074] 1372 Inf Inf Inf Inf ...
##   ..$ y.pos              : num [1:12074] 547 Inf Inf Inf Inf ...
##   ..$ identity           : num [1:12074] 434 434 434 434 434 434 434 434 434 434 ...
##   ..$ frame              : num [1:12074] 93065 93066 93067 93068 93069 ...
##   ..$ runTimelinef       : num [1:12074] 93065 93066 93067 93068 93069 ...
##   ..$ Measured_Temp_Deg_C: num [1:12074] 39.1 39.1 39.1 39.1 39.1 39.1 39.1 39.1 39.1 39.1 ...
##  $ 435:'data.frame': 12079 obs. of  9 variables:
##   ..$ maj.ax             : num [1:12079] 35.9 Inf Inf Inf Inf ...
##   ..$ angle              : num [1:12079] -0.167 Inf Inf Inf Inf ...
##   ..$ min.ax             : num [1:12079] NA NA NA NA NA NA NA NA NA NA ...
##   ..$ x.pos              : num [1:12079] 981 Inf Inf Inf Inf ...
##   ..$ y.pos              : num [1:12079] 987 Inf Inf Inf Inf ...
##   ..$ identity           : num [1:12079] 435 435 435 435 435 435 435 435 435 435 ...
##   ..$ frame              : num [1:12079] 93060 93061 93062 93063 93064 ...
##   ..$ runTimelinef       : num [1:12079] 93060 93061 93062 93063 93064 ...
##   ..$ Measured_Temp_Deg_C: num [1:12079] 39.1 39.1 39.1 39.1 39.1 39.1 39.1 39.1 39.1 39.1 ...
```

### 3 Cleaning the data

#### 3.1 Remove infinite values

##### 3.1.1 Specify the filter to detect infinite values on “x.pos”

```
filter.InfX <-
  MoveR::filterFunc(
    trackDat,
    toFilter = "x.pos",
    customFunc = function(x)
      is.infinite(x)
  )
```

*NB: it is also possible to group two or more filters by using the mergeFilter function, for instance, by merging the result of several condition tests, here is the detection of infinite value in “x.pos” and “y.pos”.*

##### 3.1.2 Specify the filter to detect infinite values on “y.pos”

```
filter.InfY <-
  MoveR::filterFunc(
    trackDat,
    toFilter = "y.pos",
    customFunc = function(x)
      is.infinite(x)
  )
```

##### 3.1.3 Merge the previously specified filters

```
filter.Inf <-
  MoveR::mergeFilters(filters = list(filter.InfX, filter.InfY),
                      cond = TRUE)
```

##### 3.1.4 Filter infinite values according to the previously specified filters

*NB: here we are also removing the tracklets that are shorter than 25 frames (1 second - the frame rate) using the minDur argument.*

```
trackDat.Infilt <-
  MoveR::filterTracklets(trackDat,
                         filter = filter.Inf,
                         splitCond = TRUE,
                         minDur = frameRate)
```

##### 3.1.5 Display the summary of the filtering process

```
str(trackDat.Infilt[[1]])
```

```
## List of 8
##  $ Tracknb_before_filter         : int 88
##  $ Tracknb_after_filter          : int 165583
##  $ Tracknb_after_minDur          : int 6724
##  $ TotTrackDuration_before_filter: int 4716398
##  $ TotTrackDuration_after_filter : int 1664533
##  $ TotTrackDuration_after_minDur : int 1177729
##  $ %Data_kept_after_filter       : num 35.3
##  $ %Data_kept_after_minDur       : num 25
```

#### 3.2 Remove individuals detected outside the arena

##### 3.2.1 Import the location of the arena edge

*NB: here we are also retrieving the location of the arena edge from a distance matrix generated using color thresholding in imageJ. As color thresholding returns a matrix with image resolution (x in rows and y in columns) and increasing distance to the arena’s edge, here the edge corresponds to the lower value of the distance matrix (i.e., 1).*

```
edge <-
  MoveR::locROI(Path2Data[[2]],
                edgeCrit = 1,
                xy = 1,
                order = T) # here specifying the order argument as TRUE ensures that the points delimiting the arena edge are sorted clockwise.

# it is then easy to draw the arena edge
plot(
  NULL,
  xlim = c(0, max(edge[, "x.pos"])),
  ylim = c(0, max(edge[, "y.pos"])),
  xlab = "Video width (pixels)",
  ylab = "Video height (pixels)",
  main = "Edge and centre of the arena",
  las = 1,
  cex.main = 0.9
)
graphics::polygon(
  x = edge[["x.pos"]],
  edge[["y.pos"]],
  lty = 2,
  col = adjustcolor("firebrick", alpha = 0.2)
)

# As well as the center of the arena
center <-
  data.frame(x.pos = mean(edge[["x.pos"]]), y.pos = mean(edge[["y.pos"]]))
graphics::points(
  x = center[["x.pos"]],
  y = center[["y.pos"]],
  col = "black",
  pch = 3,
  cex = 1
)
```

Figure 1: Representation of the arena edge (dashed line) and center (cross mark). The interior of the arena is colored in light-red.

##### 3.2.2 Draw the tracklets and add the contour of the arena

```
# draw the tracklet and the arena edges 
MoveR::drawTracklets(
  trackDat.Infilt[[2]],
  timeCol = "runTimelinef",
  add2It = list(graphics::polygon(x = edge$x.pos, y = edge$y.pos))
)
```

Figure 2: individuals trajectories colored according to the timeline and arena edge (black line).

##### 3.2.3 Assign The ROI (here the arena) to the individuals’ positions

```
trackDat2 <-
  MoveR::analyseTracklets(
    trackDat.Infilt[[2]],
    customFunc = list(
      Arena = function(x)
        MoveR::assignROI(
          x,
          ROIs = edge,
          edgeInclude = F,
          order = T
        )
    )
  )
```

##### 3.2.4 Specify the filter to detect individuals outside the arena

```
filter.out <-
  MoveR::filterFunc(
    trackDat2,
    toFilter = "Arena",
    customFunc = function(x)
      x != "ROI_1"
  )
```

##### 3.2.5 Filter individuals outside the arena according to the previously specified filters

*NB: here we are also removing the tracklets that are shorter than 25 frames (1 second - the frame rate) using the minDur argument.*

```
trackDat.borderfilt <-
  MoveR::filterTracklets(trackDat2,
                         filter.out,
                         splitCond = TRUE,
                         minDur = frameRate)

# rename the cleaned dataset for further use
trackDat3 <- trackDat.borderfilt[[2]]

# display the information about the filtering process (see ??filterTracklets())
str(trackDat.borderfilt[[1]])
```

```
## List of 8
##  $ Tracknb_before_filter         : int 6724
##  $ Tracknb_after_filter          : int 6772
##  $ Tracknb_after_minDur          : num 6707
##  $ TotTrackDuration_before_filter: int 1177729
##  $ TotTrackDuration_after_filter : int 1177327
##  $ TotTrackDuration_after_minDur : int 1176751
##  $ %Data_kept_after_filter       : num 100
##  $ %Data_kept_after_minDur       : num 99.9
```

### 4 Identify the tracklets within an ROI

#### 4.1 compute the radius of the arena in pixels

```
arenaRadius <- mean(MoveR::dist2Pt(edge, center)[[1]])
```

#### 4.2 Generate ROIs coordinates

*NB: here we are generating (circular) ROI edge coordinates with increasing distance to the center of the arena.*

```
ROIs <- MoveR::circles(
  x = rep(mean(edge[["x.pos"]]), 4),
  y = rep(mean(edge[["y.pos"]]), 4),
  radius = seq(
    from = 0,
    to = arenaRadius,
    length.out = 5
  )[2:5],
  draw = F
)
```

#### 4.3 Assign ROIs’ identity to individual’s position

```
trackDat3 <- MoveR::analyseTracklets(
  trackDat3,
  customFunc = list(
    ROIid = function(x)
      MoveR::assignROI(x, ROIs = ROIs)
  )
)
```

#### 4.4 Display colored tracklet according to the area where they are located

```
MoveR::drawTracklets(trackDat3,
                     colId = "ROIid",
                     colGrad = viridis::viridis(5),
                     legend.title = "ROI location",
                     cex.axis = 1.2,
                     cex.lab = 1.4,
                     cex.leg = 0.75
                     )
```

Figure 3: individuals trajectories colored according to their location among four circular R.O.I.s. Colors indicate the ROI where individuals are detected. As larger R.O.I.s include smaller ones, individuals located at the arena’s center are considered within the R.O.I. 1, 2, 3, and 4.

### 5 Compute some metrics

#### 5.1 Compute basic descriptors

*NB: here we are first specify a batch of functions to pass to the `analyseTracklet` function to compute metrics over each tracklet.*

```
customFuncList = list(
  # basic descriptors
  ## speed 
  speed = function(x)
    MoveR::speed(x,
                 timeCol = "runTimelinef",
                 scale = 1),
  ## turning angles
  turnAngle = function(x)
    MoveR::turnAngle(
      x,
      unit = "radians",
      timeCol = "runTimelinef",
      scale = 1
    ),
  ## sinuosity
  sinuosity = function(x)
    MoveR::sinuosity(x, 
                     timeCol = "runTimelinef", 
                     scale = 1),
  # Smooth
  ## turning angles variance
  slideVarAngle = function (y)
    MoveR::slidWindow(circular::circular(y$turnAngle,
                                         type = "angle",
                                         units = "radians"),
                      Tstep = 10, 
                      statistic = "circular.var",
                      na.rm = T),
  ## speed
  slideMeanSpeed = function (y)
    MoveR::slidWindow(y$speed,
                      Tstep = 10, 
                      statistic = "mean", 
                      na.rm = T)
)
```

```
# compute the specified basic descriptors over tracklets
trackDat4 <-
  MoveR::analyseTracklets(trackDat3,
                          customFunc = customFuncList)
```

#### 5.2 Activity states: 2D non-hierarchical clustering

##### 5.2.1 Run the classification

*NB: here we used density-based clustering algorithm to classify actives and inactive states in a two dimension array; i.e., the smoothed speed (log) and the turning angle variance.*

```
trackDat4 <-
  MoveR::activity2(
    trackDat = trackDat4,
    var1 = "slideMeanSpeed",
    var2 = "slideVarAngle",
    var1T = log10,
    nbins = 100,
    eps = 0.15,
    minPts = 5
  )
```

```
## [1] "Clusters identified !"
```

##### 5.2.2 Display the resulting clusters

*NB: It is then easy to represent the cluster in a 3d plot using the `countMat` function from MoveR’s utilites and an arbitrary graphical function made to make 3d plots (e.g., plotly).*

###### 5.2.2.1 convert the data to a count matrix

```
# compute the count matrix in the 2d space with the same parameter than activity2 function
trackDatL <- MoveR::convert2List(trackDat4)
outP <- MoveR::countMat(
  x = log10(trackDatL[["slideMeanSpeed"]]),
  y = trackDatL[["slideVarAngle"]],
  groups = trackDatL[["activity2"]],
  nbins = 100,
  output = "matrix"
)

# retrieve max and min value of sqrt-transformed count over the groups for plotting
maxVal <- max(unlist(lapply(outP, function(z)
  max(sqrt(
    z
  )))))
minVal <- min(unlist(lapply(outP, function(z)
  min(sqrt(
    z
  )))))
```

###### 5.2.2.2 use plotly to display the clusters (active and inactive) with a 3d plot

```
# draw the plot using plotly (3d interactive plot)
library(plotly)
fig <-
  plotly::plot_ly(
    x =  ~ colnames(outP[[1]]),
    y =  ~ rownames(outP[[1]]),
    contours = list(
      z = list(
        show = TRUE,
        start = round(minVal, -2),
        project = list(z = TRUE),
        end = round(maxVal, -2),
        size = max(maxVal) / 10,
        color = "white"
      )
    )
  )
# add inactive layer
fig <- plotly::add_surface(
  p = fig,
  z = sqrt(outP[[1]]),
  opacity = 0.8,
  colorscale = "Hot",
  cmin = min(sqrt(outP[[1]])),
  cmax = max(sqrt(outP[[1]])),
  colorbar = list(title = "inactive\ncounts (sqrt)")
)
# add active layer
fig <- plotly::add_surface(
  p = fig,
  z = sqrt(outP[[2]]),
  opacity = 1,
  colorscale = list(
    c(0, 0.25, 1),
    c("rgb(20,20,20)", "rgb(58,139,44)", "rgb(234,239,226)")
  ),
  colorbar = list(title = "active\ncounts (sqrt)")
)
fig <- plotly::layout(
  fig,
  title = '3D density plot of activity states',
  scene1 = list(
    xaxis = list(title = "Speed (log10)"),
    yaxis = list(title = "Angle variance"),
    zaxis = list(title = "counts (sqrt)")
  ),
  font = list(
    family = "Arial",
    size = 14,
    color = "black"
  )
)
fig
```

Figure 4: 3d density map of the active and inactive states according to the non-hierarchical classification (density based clustering) performed on smoothed individuals speed (log-transformed) and turning angle variance. here the distribution of active and inactive state are represented as a gradient of dark-green to white and dark-red to white according to the increasing number of counts.

*NB: Accordingly, it is also easy to split the dataset according to active and inactives moments and then display both active and inactive tracklets parts.*

###### 5.2.2.3 split the dataset and display active and inactive moment in separate plots

```
# display the inactive and active trajectories
## retrieve only inactive moments
activFilt <- MoveR::filterFunc(
  trackDat4,
  toFilter = "activity2",
  customFunc = function(x)
    x == 0
)

activityData <-
  stats::setNames(lapply(list(TRUE, FALSE), function(x)
    MoveR::filterTracklets(trackDat4,
                           filter = activFilt,
                           splitCond = x)),
    c("active", "inactive"))

## display a part of inactive and active moments to have an quick overview of tracklets characteristics (only the first 2000 tracklets)
par(mfrow = c(2, 1))
for (i in 1:2) {
  MoveR::drawTracklets(
    activityData[[i]][[2]],
    selTrack = ceiling(seq(1, length(activityData[[i]][[2]]), length.out = 2000)),
    timeCol = "runTimelinef",
    main = ifelse(i==1, "Actives", "Inactives"),
    add2It = list(graphics::polygon(
      x = edge$x.pos, y = edge$y.pos
    ))
  )
}
```

Figure 5: Representation of a sample of active (upper panel) and inactive (lower panel) parts of individual trajectories (i.e., 2000 tracklets).

#### 5.3 Compute and draw temporal trends

##### 5.3.1 Temporal trends

*NB: As to compute basic descriptors over tracklet, here we are first specify a batch of functions to pass to the `temporalTrend` function to compute metrics over time.*

```
# specify a list of custom function to smooth the existing metrics over time
customFuncList <- list(
  # smooth sinuosity over time
  sinuosity =
    function(x) {
      mean(x$sinuosity, na.rm = T)
    },
  # smooth speed over time for active states only
  speed_active =
    function(x) {
      ifelse(nrow(x[which(!is.na(x$activity2) &
                            x$activity2 == 1), ]) == 0, NA,
             mean(x$slideMeanSpeed[which(!is.na(x$activity2) &
                                           x$activity2 == 1)], na.rm = T))
    },
  # smooth activity states over time
  activity =
    function(x) {
      if (nrow(x[!is.na(x$activity2), ]) == 0) {
        NA
      } else {
        nrow(x[!is.na(x$activity2) &
                 x$activity2 == 1, ]) / nrow(x[!is.na(x$activity2), ])
      }
    }
)
```

```
# compute the smoothed metrics over time
TemporalRes_wtd <- MoveR::temporalTrend(
  trackDat4,
  timeCol = "runTimelinef",
  customFunc = customFuncList,
  Tstep = 90 * 25,
  sampling = 20 * 25,
  wtd = TRUE
)

# convert the timeline into minutes for graphical output
TemporalRes_wtd <- lapply(TemporalRes_wtd, function(x) {
  runTimelineMin <- x[["runTimelinef"]] / frameRate / 60
  cbind(x, runTimelineMin)
})

# append temperature values for graphical output
TempData <-
  data.frame(runTimelinef = trackDatL[["runTimelinef"]], Measured_Temp_Deg_C = trackDatL[["Measured_Temp_Deg_C"]])
TempData <- TempData[!duplicated(TempData), ]
TemporalRes_wtd <- lapply(TemporalRes_wtd, function(x)
  merge(x,
        TempData,
        by = "runTimelinef",
        all.x = T))
```

*NB: Here we can then compute the studentized 95% confidence interval using `temporalTrend` function.*

##### 5.3.2 Studentized 95% confidence interval (CI)

```
TemporalResBOOT_wtd <-
  MoveR::temporalBoot(
    trackDat = trackDat4,
    timeCol = "runTimelinef",
    customFunc = customFuncList,
    Tstep = 90 * 25,
    sampling = 20 * 25,
    bootn = 500,
    wtd = TRUE
  )

# retrieve the CI from the output of temporalBoot
boot.ci <- TemporalResBOOT_wtd[[c(which(names(TemporalResBOOT_wtd) == "BootCiStudent"))]]
```

##### 5.3.3 Plot the trends & studentized 95% CIs

*NB: We can then display the trends and studentized 95% confidence interval over time for the three specified metrics using any graphical functions, here base R.*

```
# Display temporal trends and studentized 95% CI
## remove the NA to draw the CI as an envelope
boot.ci.NoNA <- lapply(boot.ci, function(x) na.omit(x))

## convert the timeline into minutes for grapgical output
boot.ci.NoNA <- lapply(boot.ci.NoNA, function(x) {
  runTimelineMin <- x[["runTimelinef"]]/frameRate/60
  cbind(x, runTimelineMin)
})

## set the name of variable to display them on the graph
varName <- c("Sinuosity", "Speed", "Activity")

## draw the temporal trends and studentized 95% CI
par(mfrow = c(1, 3), mar = c(5, 5, 4, 2) + 0.1)
for (p in seq_along(boot.ci.NoNA)) {
  plot(
    NULL,
    ylim = c(round(
      min(c(boot.ci.NoNA[[p]][["2.5%"]], boot.ci.NoNA[[p]][["97.5%"]]) , na.rm = T), digits = max(boot.ci.NoNA[[p]][["97.5%"]], na.rm =
                                                                                                    T)
    ),
    round(
      max(c(boot.ci.NoNA[[p]][["2.5%"]], boot.ci.NoNA[[p]][["97.5%"]]), na.rm = T), digits = max(boot.ci.NoNA[[p]][["97.5%"]], na.rm =
                                                                                                   T)
    )),
    xlim = c(min(boot.ci.NoNA[[p]][["runTimelineMin"]], na.rm = T), max(boot.ci.NoNA[[p]][["runTimelineMin"]], na.rm = T)),
    main = paste(varName[[p]], "and \nstudentized 95% CI", sep = " "),
    xlab = "Timeline (minutes)",
    ylab = "",
    las = 1,
    cex.lab = 1.8,
    cex.main = 2,
    cex.axis = 1.5
  )
  title(
    ylab = varName[[p]],
    line = 3.5,
    cex.lab = 1.8,
    family = "Arial"
  )
  plotId <- LETTERS[seq_along(boot.ci.NoNA)]
  mtext(plotId[p], side = 3, line = 2.5, adj = 0, cex = 1.5)
### add the smoothed trend 
  lines(TemporalRes_wtd[[p]][[2]] ~ TemporalRes_wtd[[p]][["runTimelineMin"]],
        col = "darkred",
        lwd = 1.8)
### add the points sampled over bootstrap
  points(
    boot.ci.NoNA[[p]][["mean"]] ~ boot.ci.NoNA[[p]][["runTimelineMin"]],
    pch = 19,
    cex = 0.25,
    col = adjustcolor("grey40", alpha = 0.5)
  )
### add the 95% CI envelope
  lines(
    boot.ci.NoNA[[p]][["97.5%"]] ~ boot.ci.NoNA[[p]][["runTimelineMin"]],
    col = adjustcolor("darkred", alpha = 0.25),
    lwd = 0.5
  )
  lines(
    boot.ci.NoNA[[p]][["2.5%"]] ~ boot.ci.NoNA[[p]][["runTimelineMin"]],
    col = adjustcolor("darkred", alpha = 0.25),
    lwd = 0.5
  )
  polygon(
    x = c(boot.ci.NoNA[[p]][["runTimelineMin"]],
          rev(boot.ci.NoNA[[p]][["runTimelineMin"]])),
    y = c(boot.ci.NoNA[[p]][["2.5%"]], rev(boot.ci.NoNA[[p]][["97.5%"]])),
    col = adjustcolor("darkred", alpha = 0.1),
    border = NA
    ,
    density = NA
  )
### add the temperature threshold of 45 degrees on the plot
  maxT <-
    TemporalRes_wtd[[p]][which(TemporalRes_wtd[[p]][["Measured_Temp_Deg_C"]] == max(TemporalRes_wtd[[p]][["Measured_Temp_Deg_C"]])), ]
  abline(v = median(maxT$runTimelineMin),
         lty = 2,
         lwd = 2)
  mtext(
    expression("45\u00B0C"),
    side = 3,
    line = -1.5,
    adj  = 0.68
  )
}
```

Figure 6: Evolution of the sinuosity (A) speed (B) and activity (C) of Trichogramma individuals (dark red line) and studentized 95% confidence interval (light red envelope) over time (expressed in frame). The dashed line indicates when the maximum temperature has been reached over the ramp: 45°C.
